# Fluoro-forest: A random forest workflow for cell type annotation in high-dimensional immunofluorescence imaging

**DOI:** 10.1101/2025.06.13.659547

**Authors:** Joshua Brand, Wei Zhang, Evie Carchman, Huy Q. Dinh

## Abstract

Cyclic immunofluorescence (IF) techniques enable deep phenotyping of cells and help quantify tissue organization at high resolution. Due to its high dimensionality, workflows typically rely on unsupervised clustering, followed by cell type annotation at a cluster level for cell type assignment. Most of these methods use marker expression averages that lack a statistical evaluation of cell type annotations, which can result in misclassification. Here, we propose a strategy through an end-to-end pipeline using a semi-supervised, random forests approach to predict cell type annotations. Our method includes cluster-based sampling for training data, cell type prediction, and downstream visualization for interpretability of cell annotation that ultimately improves classification results. We show that our workflow can annotate cells more accurately with a training set <5% of the total number of cells tested. In addition, our pipeline outputs cell type annotation probabilities and model performance metrics for users to decide if it could boost their existing clustering-based workflow results for complex IF data.

**Availability and implementation:** Fluoro-forest is freely available on github (https://github.com/Josh-Brand/Fluoro-forest). Data used within this manuscript is hosted on Dryad (DOI: 10.5061/dryad.hqbzkh1v1)

**Supplementary information:** Supplemental figures and methods are included in the submission.

## 1 Introduction

High-plex IF imaging has significantly expanded our ability to visualize and characterize cell types and subsets, offering detailed views of tissue-level biology. While these technologies have existed for years^1–3^, their accessibility through commercial platforms has significantly improved. Imaging-based antibody panels can now exceed 50 markers and are more widely used, such as with Codex^4^, but open-source tools to accurately label cell types are still needed. Additionally, these datasets may span tens to hundreds of thousands of cells, making conventionally supervised or threshold-based approaches that use complex combinations of markers for cell type annotation infeasible. Open-source software, such as ImageJ^5^ and QuPath^6^ are excellent tools with extended functionality through plugins for cell segmentation, feature calculations, and cell annotation. QuPath’s classification algorithms include random forests, however it predicts marker positivity sequentially and does not generate composite models. This results in defining complicated prediction scenarios that a user would need to predefine, rather than generating a single classification probability offered through a multiclass method.

Other machine learning approaches include a probabilistic Bayesian framework (Celesta)^7^ and feed-forward neural networks (MAPS^8^, CellSighter^9^) for cell type classification. Celesta uses a design matrix to guide cell type annotation and does not require training data. This approach assumes some markers will not overlap but may intuitively struggle where fluorescence bleed over from dense areas in tissue exists. MAPS and CellSighter require large amounts of training data from thousands to tens of thousands of cells which are supplied through other cell type annotation methods. While clustering is commonly applied for other single-cell omics technologies, its use remains challenging in imaging data^10^. Aggregate expression profiles of clusters are difficult to interpret in cell type assignment due to background signal in imaging data, non-normal data distributions, and varying cluster resolutions that may mask rare cell types while over-clustering others. Evaluation of clustering-based and other cell-type annotation methods still requires having a well-labeled training dataset, using expert knowledge, which remains a consistent bottleneck. Our approach shows that small training datasets (200-300 cells) of a typical 2,000-10,000 cell tissue core data, with diverse representation of cell clusters, may improve cell type annotation results significantly.

## 2 Results

We developed Fluoro-forest, an end-to-end Python pipeline for cell type annotation and prediction using user-defined protein markers. Our pipeline (**Fig. 1A, Suppl. Fig. 1**) begins with cell segmentation using existing methods, such as StarDist^11^ or CellPose^12^ to identify cell boundaries, followed by pre-processing procedures including log-normalization and z-scaling within each marker (**Suppl. Methods**). The images (one slice per marker), tabular expression data, and segmentations are inputs for a Python class for data visualization, cell sampling for training data, annotation, and cell type prediction. In our workflow, the user is prompted to select which markers (by default, all) to use in dimension reduction with principal components analysis (PCA) for visualization. We support unsupervised clustering methods (k-means, Louvain, Leiden, or Gaussian mixture models) to initialize the sampling procedure to ensure a diverse training dataset across all cell lineages. The sampled cells will be annotated by users using a multichannel viewing window and serve as training data for a random forest model to predict the remaining unannotated cells. We include optional settings such as imbalanced random forests or synthetic sampling (SMOTE^13^) to improve classification accuracy in cases where the majority cell type dominates the training data. Finally, we provide cross-validation performance metrics (precision, recall, and F1 scores) output per cell type to guide model tuning.

**Figure 1.**
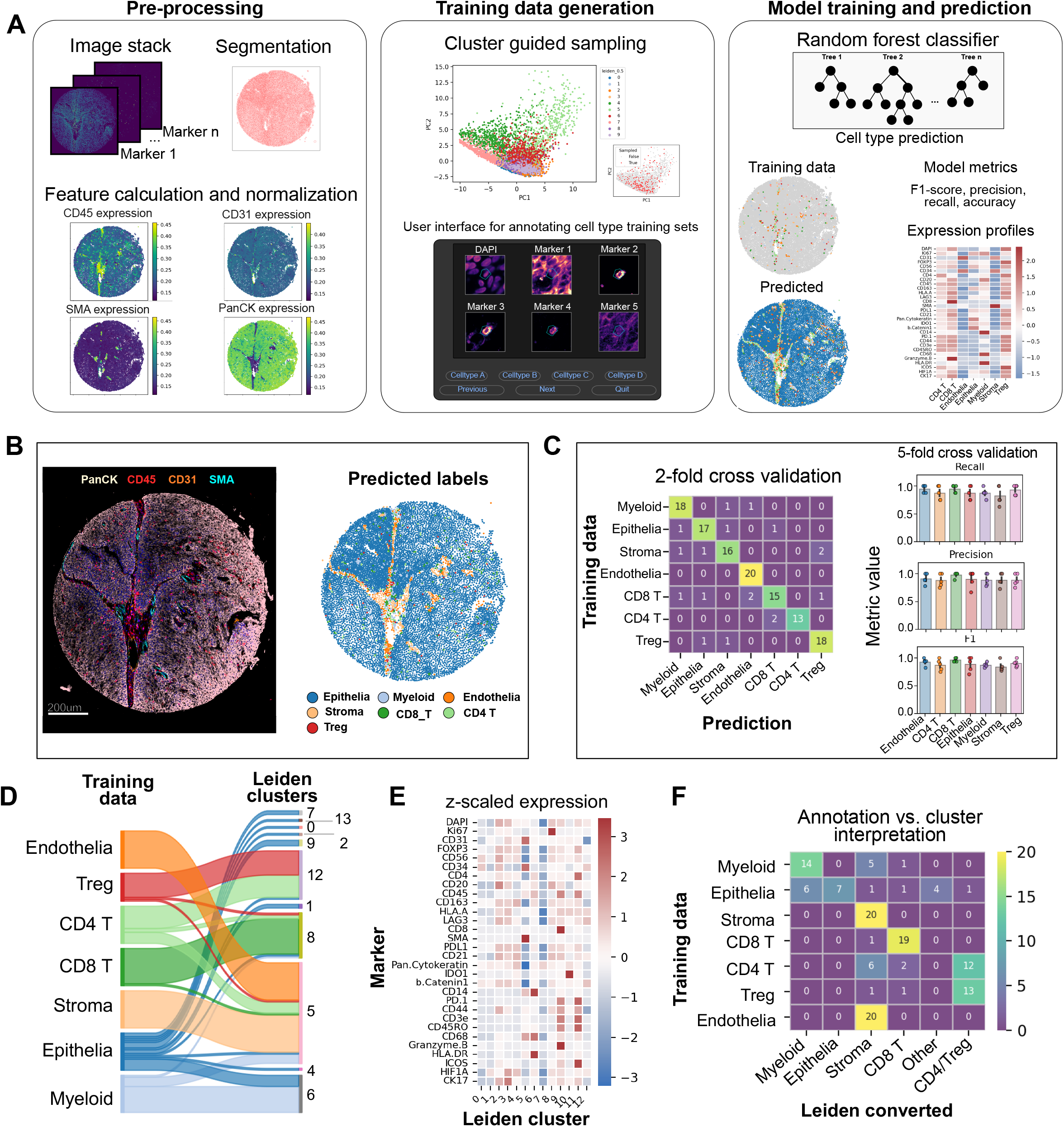
Fluoro-forest: A semi-supervised cell type annotation method using random forests for high-plex immunofluorescence data. **A**. Our workflow converts images to tabular data, guides cell annotation to train random forest models for predicting unseen data. **B**. Lineage markers for epithelia (PanCK), endothelia (CD31), smooth muscle/stroma (SMA), and immune cells (CD45) supports predicted annotations. **C**. Classes were balanced prior to training and tested for model accuracy in 2 and 5-fold cross validation, achieving 87% and 90.4 +/-2.7% accuracy, respectively (mean +/-s.d.). **D**. Sankey diagram of annotated cell types and their leiden-based clustering. **E**. Z-scored expression values for each leiden cluster across all markers. **F**. Expression guided annotation from clustering was unable to split CD4+ T cells from Tregulatory cells, and stroma from endothelia.

To illustrate the workflow, we applied our pipeline to PhenoCycler / CODEX data from two in-house anal dysplasia biopsies with an immune-focused protein panel (N = 30 markers) (**Fig. 1B, Suppl. Fig. 2A**). We identified cell types in these datasets – including tumor/epithelial cells, stroma, endothelial cells, myeloid, CD4 and CD8 T cells, B cells, and Tregs based on their representative markers (**Suppl. Methods**). Lineage marker expression well-supported our final cell type predictions in both cores (**Fig. 1B, Suppl. Fig. 2A**). Five-fold cross validation showed high accuracy (90.4 +/-2.7% and 89.1 +/-3.5% respectively, mean +/-s.d.) across our 2 biopsies (**Fig. 1C, Suppl. Fig 3B**) and demonstrate the model’s ability to identify true classes while minimizing false positives and false negatives, achieving F1 scores of 0.75 or higher. We compared our pipeline to annotations derived from a Leiden-based clustering workflow, showing lack of purity in cell type annotations from Leiden clustering results, for instance cluster 5 includes mixture of endothelial, epithelial, CD4 T and stroma cells (**Fig. 1D, Suppl. Fig. 2D**), which is challenging for cell type annotation based on average marker expression (**Fig. 1E-F, Suppl. Fig. 2D-E**). Specifically, we used all features to over-cluster the data, which is recommended^10^ to recover less abundant cell types, such as Tregs and myeloid cells. Leiden clustering was able to identify different subsets from predominantly epithelial cells but failed to differentiate FOXP3+ T regulatory cells from other CD4+ T cells and CD31+ endothelial cells from SMA+ stromal cells (**Fig. 1F, Suppl. Fig. 2E**).

## 3 Conclusion

Semi-supervised methods offer several advantages over clustering for cell type annotation in IF data. We see improved accuracy and generate interpretable probability outputs from trained models, which are not found in most clustering approaches. Our study shows that sampling approximately 20 cells per class was sufficient for high-accuracy predictions. Despite these improvements, limitations exist. Training data, and therefore validation during data splits, are represented by cells that we can confidently annotate, but acknowledge there are areas in tissue where the high cell density prevents a confident annotation. We expect the use of probability thresholds from the model can be applied for the conservative estimation of cell types in tissue where this exists. Finally, we expect these models may not generalize well, seeing a significant loss in accuracy (<70%) (**Suppl. Fig. 3A-C**) when training on one core and predicting the other. When training data are generated from both cores, there is still a loss in performance, with ∼88% accuracy in our cross-validation. We expect batch variation and donor heterogeneity to be challenges that could be overcome with additional feature engineering. However, our method offers a valuable, streamlined user interface appropriate for high-resolution and high-plex cell annotation.

## Author contributions

Conceptualization, J.B. H.Q.D., Data analysis, J.B., Data generation, E.C., W.Z., Writing J.B., H.Q.D., Review, J.B., E.C., W.Z., H.Q.D

Corresponding authors: J.B, H.Q.D.

## Funding

This work was supported by the NIH: NCI 3P30CA014520-45S1.

---

**Supplementary Figure 1.**
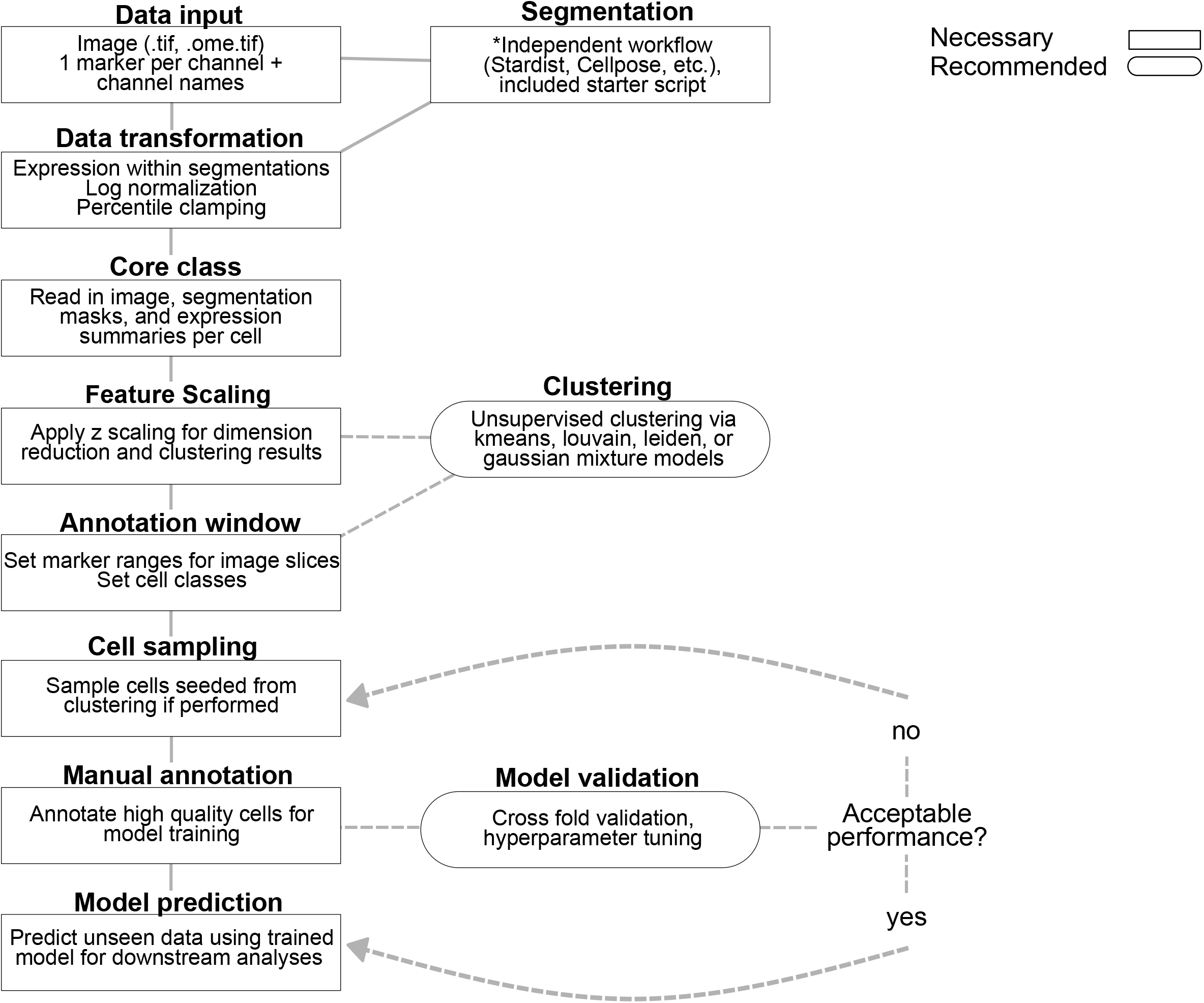
Recommended fluoro-forest workflow. Our workflow uses existing tools for cell segmentation, feature summaries, and clustering to guide cell sampling. Sampled data is shown to the user for manual annotation which trains a random forest classifier. Built in cross-fold validation metrics can guide parameter tuning, ideal models (balanced/iimbalanced), or suggest if if more samples are needed, Last, the final model using all training data can be used to predict unseen cells for downstream visualizations and analyses.

**Supplementary Fig. 2.**
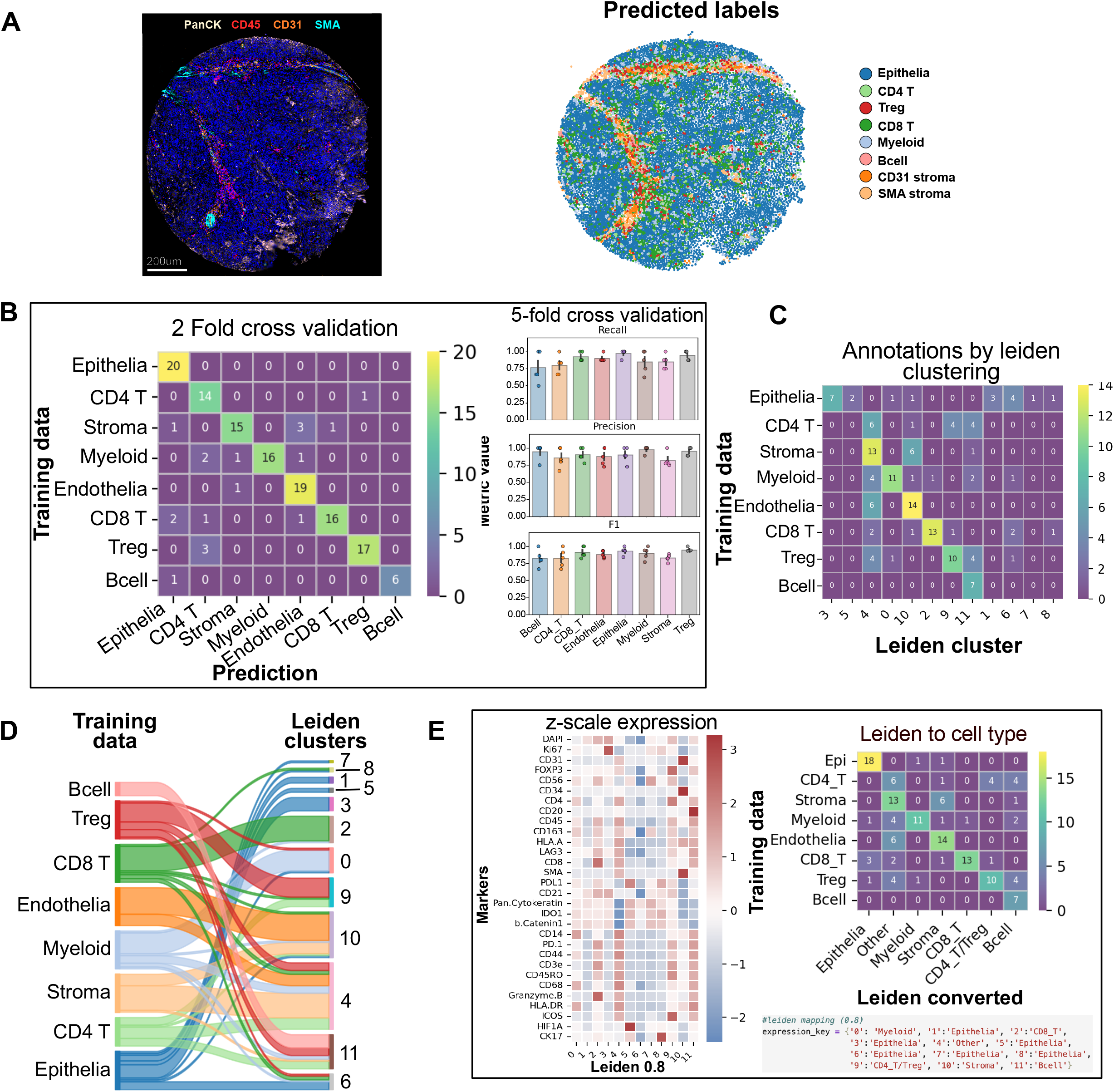
Second biopsy prediction and evaluation. **A**. Lineage markers for epithelia, stroma, endothelia, and pan immune cells next to random forest predicted cell labeled results. **B**. Random forest models were compared via 2-and 5-fold cross validation with model accuracies of 87.0% and 89.1% +/-3.5%, respectively (mean +/-s.d.). **C**. Leiden--based clustering was applied using all scaled features, showing separation of epithelia in several clusters. Clustering failed to separate stroma and endothelia, as well as CD4+ T cells and Tregs. **D**. Sankey diagram representing the annotated cell types and their representation in leiden-based clustering. **E**. Z-scored expression values for each cluster across all markers were used to annotate cell types as in a typical workflow, resulting in incomplete separation of cell types.

**Supplementary Fig. 3.**
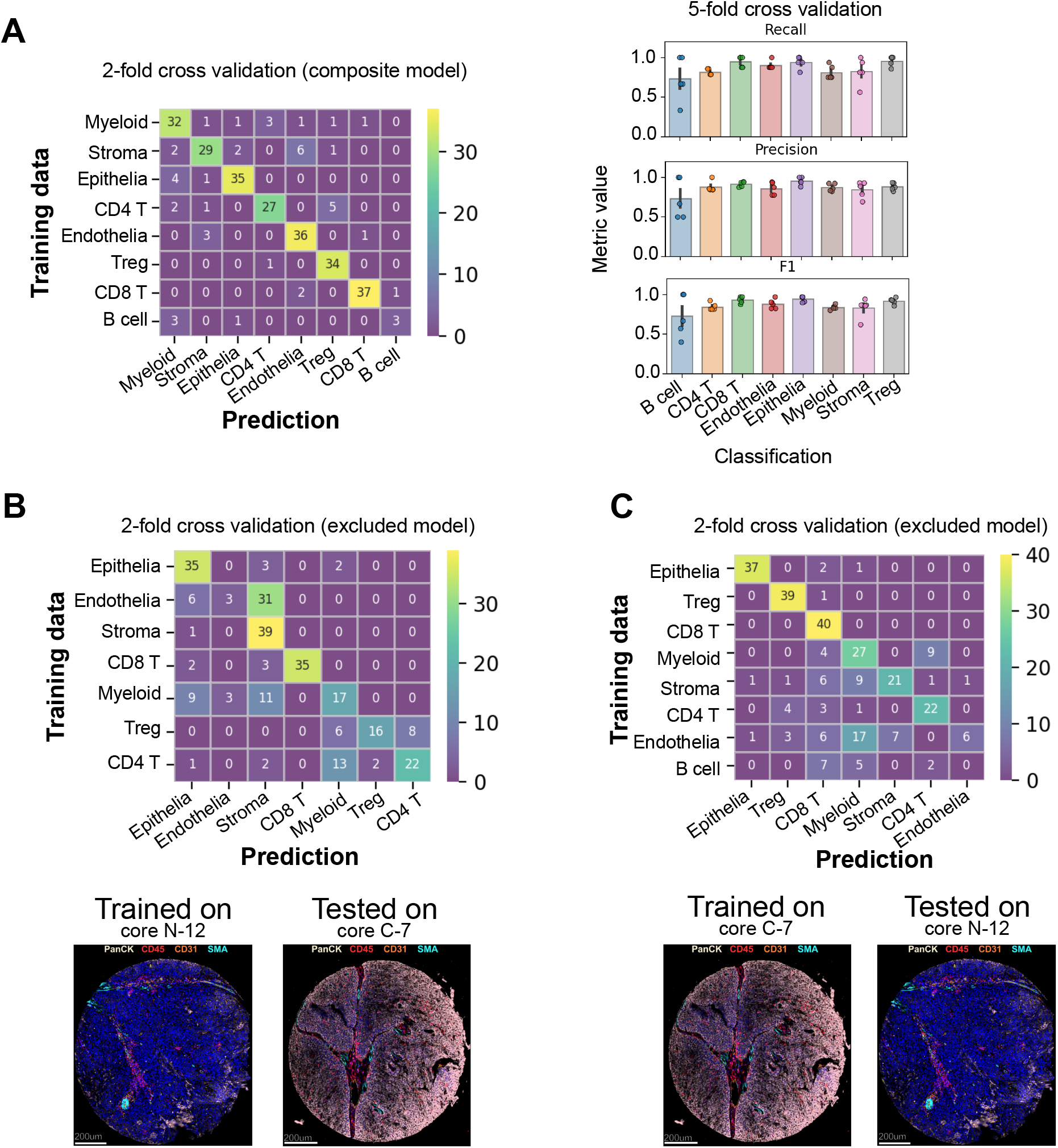
Composite vs. exclusive models shows loss in accuracy. **A**. Random forest trained on data from both cores to create a composite model, achieving 84% and 88.10 +/-2.96% accuracy, respectively (mean +/-s.d.). **B**. Classification results from training on core N-12 and testing on core C-7, 61.85% accuracy. **C**. Classification results from training on core C-7 and testing on core N-12, 67.61% accuracy.

## Notes

https://doi.org/10.5061/dryad.hqbzkh1v1

### Competing Interest Statement

The authors have declared no competing interest.

