## Supplemental Methods for "Fluoro-forest: A random forest workflow for cell type annotation in high-dimensional immunofluorescence imaging"

**Data and code access**

PhenoCycler / Codex data access for the 2 cores used in this manuscript are provided through Dryad (DOI: [10.5061/dryad.hqbzkh1v1](https://datadryad.org/submission/10.5061/dryad.hqbzkh1v1)). This submission includes images, segmentation results after applying StarDist^1^, and QC’d expression values that are used for random forest training and prediction.

Fluoro-forest is freely available on GitHub (<https://github.com/Josh-Brand/Fluoro-forest>).

Our pipeline is provided through Jupyter Notebooks found at <https://github.com/Josh-Brand/Fluoro-forest>, and includes .yml files for environment setup, relying on standardized packages – scipy, numpy, matplotlib, sklearn, seaborn for data processing, visualization, and analysis. Our GitHub readme contains details on where to store the image files (N-7.ome.tif, C-12.ome.tif) and expression ‘_expression.tsv’ files within the directory to reproduce our examples.

To reproduce the results, a .yml file is provided. After downloading the source code from GitHub, the images and expression summaries from dryad create a new virtual environment:

*conda env create -f cell_annotation.yml*

Enter into the project directory and run jupyter notebook / jupyter lab to access your notebooks. Proceed with running core-c7.ipynb, which walks through each step of data loading, clustering, sampling, annotation, and predictions.

**Data generation**

Selected samples from anal precancers and cancers were used to create a tumor microarray (TMA) for spatial phenotyping analysis using the Akoya Phenocycler Fusion (formerly known as CODEX). CODEX data were generated at The Bursky Center for Human Immunology and Immunotherapy Programs, Washington University School of Medicine. This approach uses tissue based cyclic immunofluorescence for highly multiplexed immunofluorescence imagining on FFPE specimens from glass slides. Data shown within this manuscript were 2, 2mm core biopsies sourced from at the University of Wisconsin-Madison using a custom panel (below). Final stitched images for the cores used here are available via Dryad in as .ome.tif files which were used for cell annotation.

30 maker panel: each image slice corresponding to the marker in the order shown

DAPI, Ki67, CD31, FOXP3 ,CD56	**1 - 5,**

CD34, CD4, CD20, CD45, CD163	**6 - 10,**

HLA-A, LAG3, CD8, SMA, PDL1	**11 - 15,**

CD21, PanCK, IDO1, bCat1, CD14	**16 - 20,**

PD-1, CD44, CD3e, CD45RO, CD68	**21 - 25,**

GZMB, HLA-DR, ICOS, HIF1A, CK17	**26 - 30**

**Images and figures**

Images presented in this paper were generated in QuPath^3^ (v0.5.0) and pseudo-colored to show marker localization for DAPI, PanCK, SMA, CD31, and CD45.

**Cell segmentation**

Our workflow assumes cell segmentation has been performed on a multiplex IF image, where each image slice is an independent marker (**Methods Figure 1)**. The segmentation boundaries are converted to polygons and visualized during the cell annotation procedures.

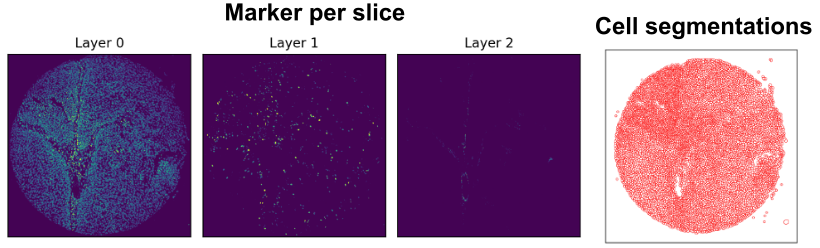

**Methods Figure 1.** Expected image layout requires each marker to be unique for each image to calculate expression summaries for each marker independently after segmentation.

In our examples, we applied a StarDist^1^, a deep learning-based segmentation method using their existing nuclear segmentation model (dsb2018_heavy_agument.pb) provided via the QuPath plugin (https://github.com/qupath/qupath-extension-stardist). Channel values were normalized to 2-98 percentiles and a pixel size of 0.5 and cell expansion of 5 was applied. Thresholds for detection using StarDist were most sensitive to changing segmentation results and therefore tested at various levels and confirmed visually before exporting the data.

The segmentation results were saved out as a .geojson file using QuPath. The .json structures the cell ids and coordinates which are parsed using utility functions before being read into into a python class for cell annotation, model training, and prediction **(Methods Figure 2**). The cell segmentations were used to summarize the expression of all markers creating features for cytoplasmic, membrane, nuclear, and cell regions as provided through QuPath’s ‘Measurement Export’ tool. Our downstream analysis only considers features calculated over the entire segmentation region (suffix with ‘_Cell_Mean’) other than DAPI, which was excluded in cell type prediction.

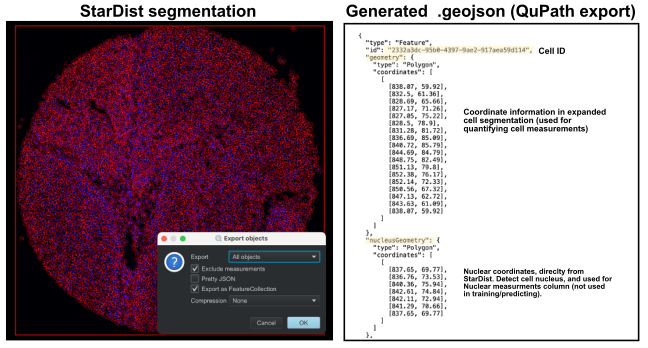

**Methods Figure 2.** QuPath segmentation after running StarDist generates a .geomjson file, which is parsed for individual cells and used to link expression data back to cell id and its location in tissue.

Note:

Our code base provides a small example of how to run segmentation exclusively in python to help new users, through an alternative cell segmentation workflow via CellPose^4^, but requires a separate python environment with .yaml provided.

*conda env create -f cellpose.yaml*

After creating this environment activate it and start with the ‘segmentation_example.ipynb’ for a step-by-step example of cell segmentation. From this environment one of the cores is read in and cropped to generate a small data example. CellPose runs its internal nuclear model for segmentation and expression results are averaged over the entire cell, which are visualized to confirm its matching expression location in tissue.

**Data transformation**

The features calculated after segmentation were exported from QuPath, along with cell ID’s, and the centroids of cells (x, y spatial location). Each core was treated independently in QC analysis, which clamped low values to 1% of the percentile value of a given marker and high values to 99%. This step is similar to existing workflows^5^ and prevents the influence of outliers on mean expression. Next, we applied a log2 transform on the percentile clamped data with a pseudocount of 0.01 to prevent log operations on intensity values of 0. Log normalized intensity was used directly for random forest training and prediction but are z-scaled when applying clustering and principal components analysis.

QC was performed to remove cells with poor quality segmentations, detected by low DAPI stain and under-stained cells. The QC’d data was read into a python notebook along with the image file (.tif), segmentation file (.geomjson from StarDist) for interactive cell annotation.

**Clustering and cell sampling**

Clustering via kmeans, Leiden, Louvain, and gaussian mixture models, are options provided to the user. Each can be calculated using all features or a subset of features of the users choosing. In our examples we applied Leiden clustering using z-scaled, cell features to improve sampling. This approach ensures sampling diversity and reduces overrepresented cell types during sampling. In cases where immune cells are expected to be more diverse (immune cells) we perform broad clustering using lineage markers and subscluster at higher resolutions.

Sampling procedures can be performed manually – through thresholding of markers to enrich for cell types. The workflow applied here was first performing a lineage-based clustering from SoftMax expression of lineage markers (PanCK, CD31, SMA, and CD45) and performing kmeans, where k = 4. Later we applied Leiden clustering within each lineage level cluster to control cell type resolutions for harder to sample cell types or where more diversity is expected to exist. We oversample within each cluster to generate a diverse training queue and annotate high quality cells with clear marker expression and minimal cell overlap. For our data we selected markers to view and annotated according to **Methods Table 1**.

**Methods Table 1.** Marker combinations for cell type annotation.

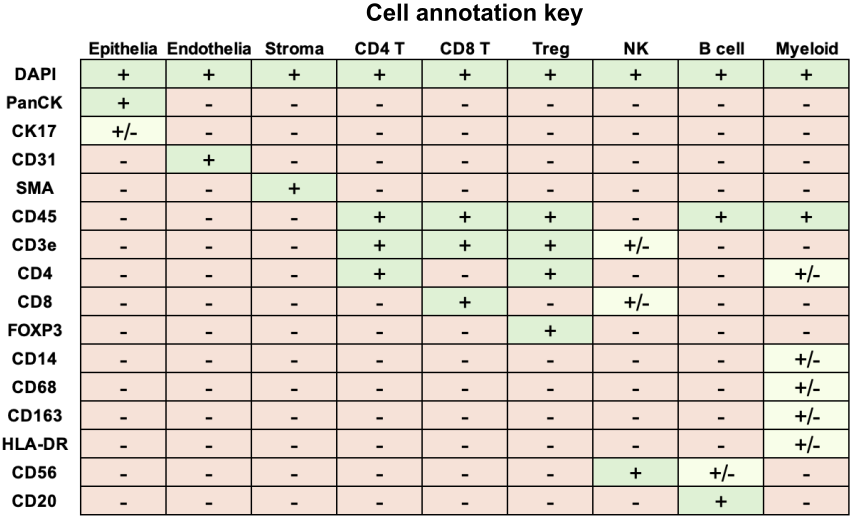

During annotation the samples selected based on clustering or any other means are provided to the user one at a time. A multi-channel viewing window is displayed with user-defined markers (**Methods Figure 3**) and annotation buttons allowing to annotate the cell type, skip to another cell, or exit the application. Upon quitting or finalizing annotations, the cell ids and their annotations are saved to the python class for later use in training and prediction.

**Cluster annotation from average expression**

Expression information was visualized using heatmaps by averaging the log-normalized expression of all values within a given annotation or cluster. These values were z-scored across annotations to demonstrate intensity relative to other groups for each marker. We used knowledge of combination and exclusive markers to attempt accurate classification with expression summaries, as is done in common workflows.

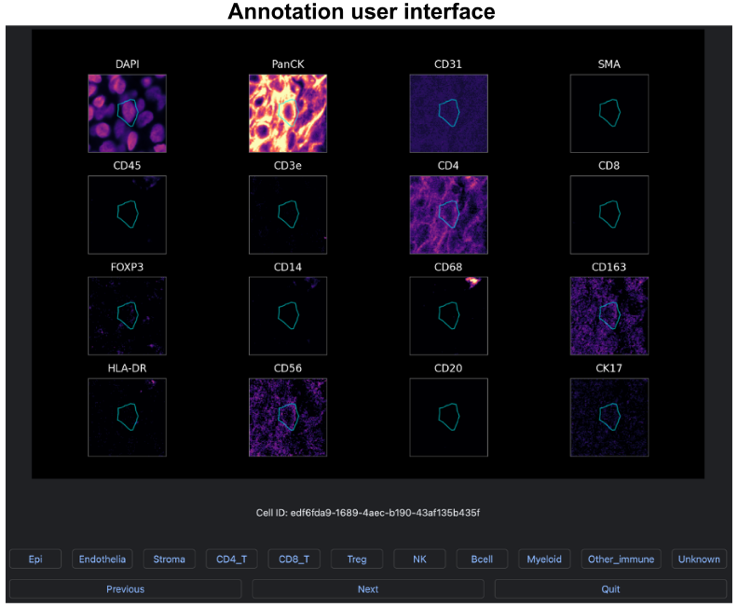

**Methods Figure 3.** Segmentations processed from .geomjson files are used to find individual cells in the image and are centered within each window. The cyan polygon represents the cell type to be annotated based visualized across channel markers in the dataset.

**Random forest training, prediction, and performance evaluation**

Cells were annotated across each core individually, guided by the clustering steps prior. Annotations were randomly down sampled to achieve a balanced representation of immune classes (B cells, myeloid, CD4 T, CD8 T, and Tregs) relative to epithelia, which were dominant in both cores. We down sampled to achieve approximately 40 cells per class, which was split into different cross validation (2 and 5-fold cross validation tests) schemes. 2-fold cross validation was used to visualize where misclassifications were occurring, while 5-fold cross validation was used to estimate model metrics such as precision: TP/(TP+FP), recall: TP/(TP+FN), and f1 scores: (2*precision*recall)/(precision + recall) for each. (TP =True positive, FP = False positive, FN = False negative).

The final predicted annotations were derived from a full model, which used the ~40 cells per class to predict the remaining unseen cells in our data. Last, we checked the performance of models trained on one core exclusively, then tested on the other or a composite model where cells from both cores were trained. The significant loss in accuracy when comparing the exclusive models suggest that data should be sampled across all cores where available for improved cell typing. Although the composite model performed with ~85% accuracy, the exclusive models support there are likely batch effects preventing high-quality, generalizable models without additional feature engineering
